# Contrasting inflammatory signatures in peripheral blood and bronchoalveolar cells reveal compartment-specific effects of HIV infection

**DOI:** 10.1101/804229

**Authors:** Daniel M. Muema, Maphe Mthembu, Abigail Schiff, Urisha Singh, Bj□rn Corleis, Thierry Bassett, Sipho S. Rasehlo, Kennedy Nyamande, Dilshaad Fakey Khan, Priya Maharaj, Mohammed Mitha, Moosa Suleman, Zoey Mhlane, Taryn Naidoo, Dirhona Ramjit, Farina Karim, Douglas S. Kwon, Thumbi Ndung’u, Emily B. Wong

**Author notes:** Corresponding author: Dr. Emily B Wong, Africa Health Research Institute, 719 Umbilo Road, Durban 4001, South Africa. These authors contributed equally (Daniel M. Muema, Maphe Mthembu).

## Abstract

The mechanisms by which HIV increases susceptibility to tuberculosis and other respiratory infections are incompletely understood. We used transcriptomics of paired whole bronchoalveolar lavage (BAL) fluid and peripheral blood mononuclear cells to compare the effect of HIV at the lung mucosal surface and in the peripheral blood. The large majority of HIV-induced differentially expressed genes (DEGs) were specific to either the peripheral or lung mucosa compartments (1,307/1,404, 93%). Type I interferon signaling was the dominant signature of DEGs in HIV-positive blood with a less dominant and qualitatively distinct type I interferon gene set expression pattern in HIV-positive BAL. DEGs in the HIV-positive BAL were significantly enriched for infiltration with cytotoxic CD8^+^ T cells. Higher expression of representative transcripts and proteins in BAL CD8^+^ T cells during HIV infection, including *IFNG* (IFN-γ), *GZMB* (Granzyme B) and *PDCD1* (PD-1), was confirmed by cell-subset specific transcriptional analysis and flow cytometry. Thus, we report that a whole transcriptomic approach revealed qualitatively distinct effects of HIV in blood and bronchoalveolar compartments. Further work exploring the impact of distinct type I interferon programs and CD8^+^ T cells infiltration of the lung mucosa during HIV infection may provide novel insights into HIV-induced susceptibility to respiratory pathogens.

## Introduction

HIV is a major cause of morbidity and mortality in sub-Saharan Africa. South Africa bears the highest burden globally with approximately 7.7 million people living with HIV in 2018 ^1^. Introduction of immediate antiretroviral therapy (ART) following diagnosis has greatly improved long-term outcomes and life expectancy in people living with HIV. However, 46% of people living with HIV in South Africa are still viremic either because they don’t know their HIV status or due to treatment failure ^1^.

HIV-infected viremic patients are more likely to develop active tuberculosis (TB) from either new exposure to *M. tuberculosis* (Mtb) or reactivation of a pre-existing latent Mtb infection when compared to HIV-uninfected and HIV-infected non-viremic individuals ^2, 3^. T helper 1 (Th1) polarized CD4^+^ T cells that produce interferon gamma and TNF-alpha are thought to play a significant role in controlling TB infection, and their depletion in HIV-infected individuals may contribute to the increased risk of TB disease ^4, 5^. However, the risk of developing active TB increases even before significant CD4^+^ T cell depletion, doubling within the first year of HIV infection, suggesting that other HIV-induced modulations of the immune system could contribute to the increased risk of TB in people with HIV ^6^. Indeed, HIV infection has been shown to decrease the polyfunctional cytokine production in Mtb-specific CD4^+^ T cells independent of CD4^+^ T cell depletion ^7, 8^. Additionally, HIV infection is associated with reduced phagocytic potential of alveolar macrophages, an effect that could reduce the initial innate immune barrier to TB infection ^9^.

Strategies to enhance control of Mtb in HIV-infected and uninfected persons are hindered by our limited understanding of the natural immunological control of Mtb and the mechanisms underlying progression from latent Mtb infection to disease. Hypothesis-driven targeted studies may miss important pathways that could inform the understanding of TB immunopathogenesis. In contrast, high throughput approaches have the potential to offer unbiased insights into the immune defects mediated by HIV, and thus advance the field of TB immunopathogenesis. One such approach is harnessing genome wide transcriptomic data to provide mechanistic insights into the immunologic pathways that are defective in patients who are likely to get infected with TB or progress to active disease, such as HIV-positive patients ^10-14^. The use of whole blood or unsorted peripheral blood mononuclear cells (PBMCs) in transcriptomic studies has the advantage of giving an overall picture of the immune profiles that are associated with development of TB. For instance, transcriptomic analyses of whole peripheral blood in tuberculosis has pointed to the development of a type 1 interferon signature ^10^. Notably, even though it is established that HIV disrupts lung immunity and increases risk of TB disease, these whole genome studies have not investigated the effect of HIV in the lung, the site of Mtb exposure; most of the work has been on samples from peripheral blood because they are easily accessible.

In this study, we assessed the differences in immune profiles between blood and the bronchoalveolar compartments using whole compartment and sorted CD8^+^ T cells by RNA-seq and flow cytometry. We also assessed the compartment-specific effects of HIV to comprehensively explore and describe the immunological defects that could explain the increased lung comorbidities, especially TB disease, in HIV-infected individuals. We report that while HIV induces primarily a type I interferon signature in blood, its primary signature in the bronchoalveolar compartment is an induction of a cytotoxic CD8^+^ T cell infiltrate.

## Results

### Population characteristics

We used matched blood and bronchoalveolar fluid samples that were collected from 19 HIV-negative participants (15 from the research bronchoscopy cohort and 4 from the hospital-based cohort) and 11 HIV-positive participants (8 from the research bronchoscopy cohort and 3 from the hospital-based cohort) (Table 1). The median age of all participants was 34 years and 50% of the participants were female. There were no significant differences in age or sex distribution between the HIV-negative and HIV-positive groups. All HIV-infected participants were ART-naïve. The median viral load for the HIV-infected group was 54,942 (Interquartile range [IQR]: 18,743-174,293). The median CD4+ T-cell counts for the HIV-uninfected and HIV-infected groups were 1,048 (IQR: 854-1,352) and 353 (IQR: 173-576), respectively (p<0.0001).

**Table 1:**
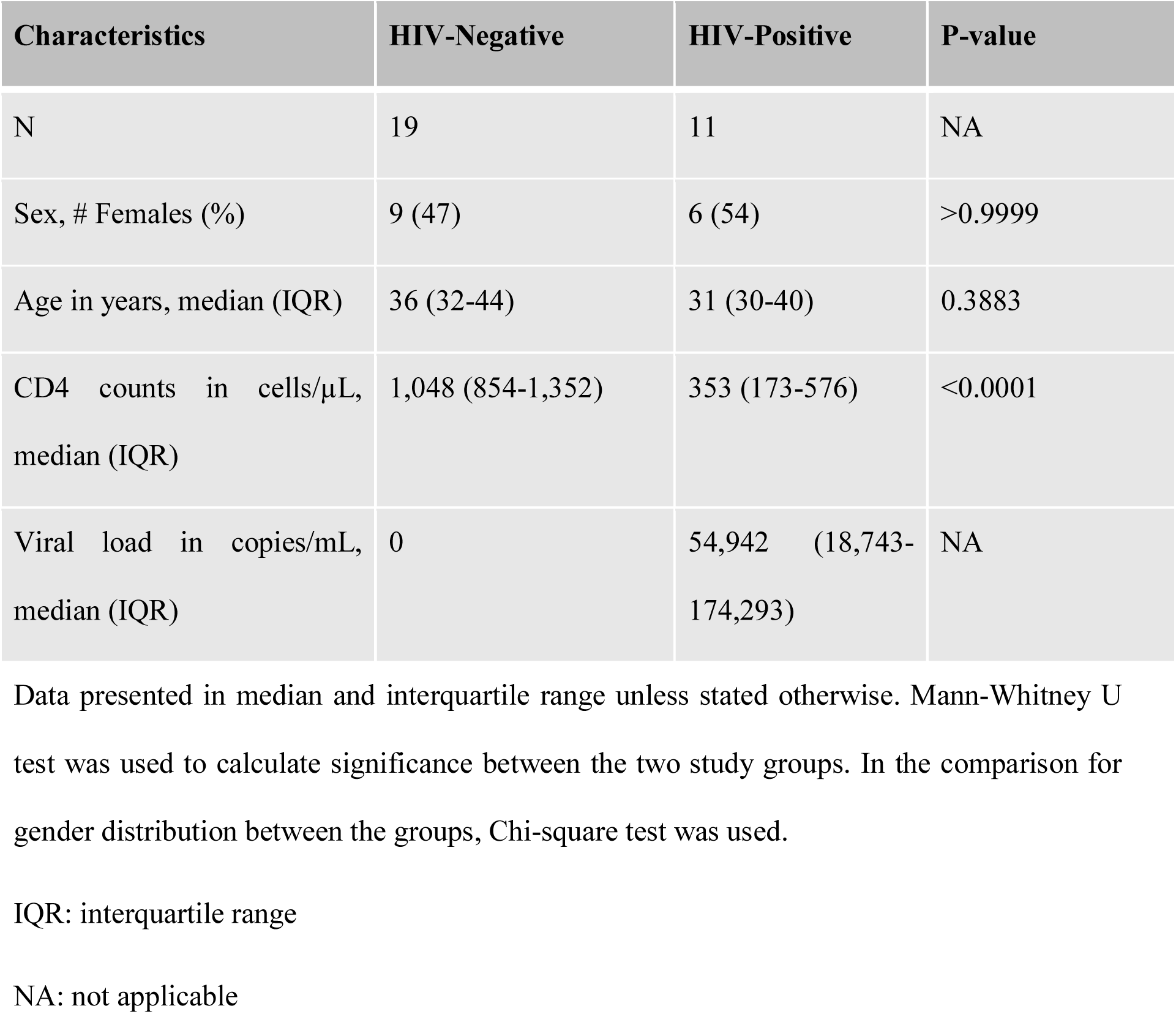
Demographic and clinical characteristics of the study population.

### Distribution of major immune cell populations in bronchoalveolar compartment and blood

To determine the HIV-specific effects on the distribution of major populations of immune cells within each anatomical compartment, we used matched samples from the bronchoalveolar compartment and blood to conduct two-way comparisons i.e. between disease states within each compartment (comparisons 1 and 2, Figure 1A) and between compartments (comparisons 3 and 4). We carried out differential cell counts to enumerate differences in distribution of key immune cell populations between the two compartments (Figure 1B). In both HIV-negative and HIV-positive individuals, distributions of immune cells were significantly different between the two compartments (Figure 1C). In both groups, alveolar macrophages were the dominant immune cell in the bronchoalveolar compartment with medians of 87.4% (Interquartile range [IQR]: 79.8-90) and 79.3 (Interquartile range [IQR]: 70.9-88) among HIV-negative and HIV-positive persons, respectively (Figure 1C and Supplementary table 1). On the other hand, lymphocytes and neutrophils were the dominant immune cells in the peripheral blood in both groups. Notably, the distributions of major immune cells were not significantly different between HIV-negative and HIV-positive participants within the compartments.

**Figure 1:**
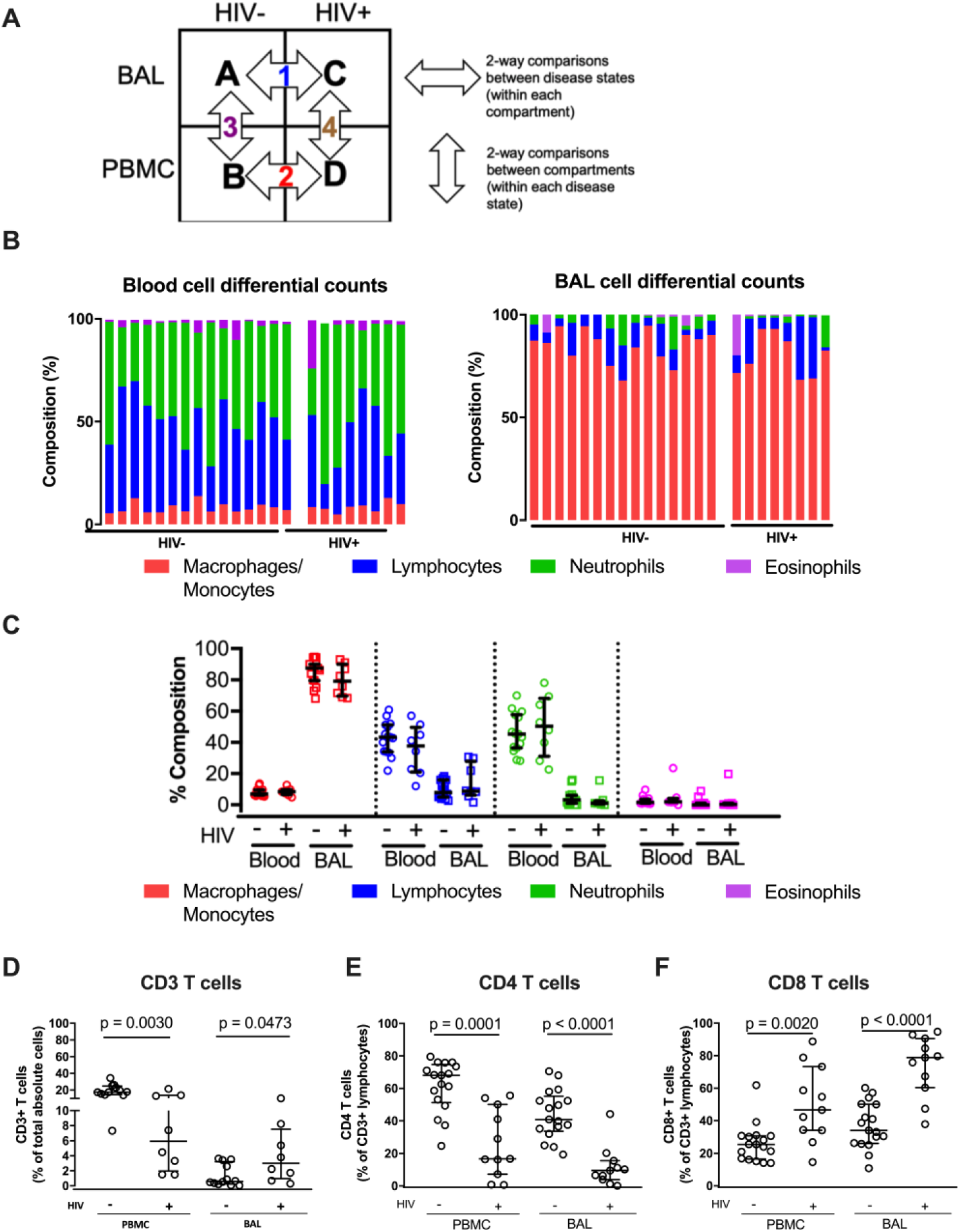
Comparisons of the immune cell sub-types between peripheral blood and bronchoalveolar lavage (BAL) cells of both HIV-negative and HIV-positive individuals. A) Two-way comparison of the study groups to understand inter-compartment differences and the impact of HIV on immune function. B) Proportion of immune cell subsets in the paired peripheral blood and bronchoalveolar cells in the HIV-negative (n=15) and HIV-positive groups (n=8). C) Quantitative comparison of panel B, showing level of significance between compartments in both HIV-negative and HIV-positive groups. D) Proportions of total absolute CD3+ T cells in BLCs and PBMCs in HIV-positive (n=8) and HIV-negative groups (n=15). E) Percentage CD4+ T cells of CD3+ T cells in BLCs and PBMCs in HIV-positive (n=11) and HIV-negative groups (n=19). F) Percentage of CD8+ T cells in BLCs and PBMCs in HIV-positive (n=11) and HIV-negative groups (n=19). The * in the figures means p value was < 0.05.

We then used flow cytometry to further assess if there were HIV-specific alterations within the lymphocyte populations. The proportions of total T cells (CD3^+^ cells) were reduced in the peripheral blood mononuclear cells (PBMCs) but increased in bronchoalveolar lavage fluid cells (BLCs) of HIV-positive patients, consistent with HIV-associated T-cell infiltration in bronchoalveolar compartment (p = 0.0030 and p=0.0473, respectively) (Figure 1D). The reduction of proportions of CD3^+^ lymphocytes in PBMCs of HIV-positive study participants was primarily due to the loss of CD4^+^ T cells as shown by the reduction in proportions of CD4^+^ T cells in PBMCs (p =0.0001). We observed similar reduction in proportions of CD4^+^ T cells in the BLCs of HIV-positive participants (p <0.0001) (Figure 1E). Notably, we observed an HIV-associated increase in proportions of CD8^+^ T cells in both PBMCs (p = 0.002) and BLCs (p = <0.0001). In further separate analyses of the research bronchoscopy cohort and the hospital-based cohort, we observed a similar HIV-associated increase in proportions of CD8^+^ T cells and a reduction in proportions of CD4^+^ T cells in the BLCs and PBMCs (Supplementary figure 4 A, B, D and E). Thus, HIV was associated with an increase in proportions of CD8^+^ T cells and a decrease in proportions of CD4^+^ T cells in both blood and bronchoalveolar compartments in multiple cohorts.

### HIV infection is associated with compartment-specific changes in the transcriptional profile in BLCs and PBMCs

To first assess transcriptome-wide differences between compartments, we used RNA-seq to determine RNA expression differences between BLCs and PBMCs in four HIV-uninfected participants and three HIV-infected participants from the hospital-based cohort. Except for one PBMCs sample, we obtained at least 1 million unique forward and reverse reads for each sample after deduplication (Supplementary figure 2).

There were 4,761 differentially expressed genes (DEGs) between the BLCs and PBMCs in either HIV-positive or HIV-negative participants. Of these, there were 4,084 DEGs between BLCs and PBMCs in the HIV-negative group, with 2,336 genes being upregulated and 1,748 genes being downregulated in BLCs when compared with PBMCs (Figure 2A). On the other hand, there were 2,186 DEGs between BLCs and PBMCs in the HIV-positive group, with 1,204 genes being upregulated and 982 genes being downregulated in BLCs when compared with PBMCs (Figure 2B). Notably, the large majority of the DEGs (69% (1,509 out of 2,186)) between compartments in the HIV-positive individuals were also differentially expressed between compartments in the HIV-negative individuals (Figure 2C).

To assess the compartment-specific effects of HIV, we then checked for differences between the HIV-negative and HIV-positive groups within each compartment. There were 774 DEGs in PBMCs between the HIV-positive and the HIV-negative groups, with 515 genes being upregulated and 259 genes being downregulated in the HIV-positive group (Figure 2D). On the other hand, there were 727 DEGs in BLCs in comparisons between the HIV-positive group and the HIV-negative group, with 540 genes being upregulated and 187 genes being downregulated in the HIV-positive group (Figure 2E). Notably, of the DEGs in either the BLCs or the PBMCs between disease states, only a very small minority (6.9%, 97 of 1401) were differentially expressed in both compartments. Thus, HIV-induced transcriptional alterations were compartment-specific (Figure 2F).

**Figure 2:**
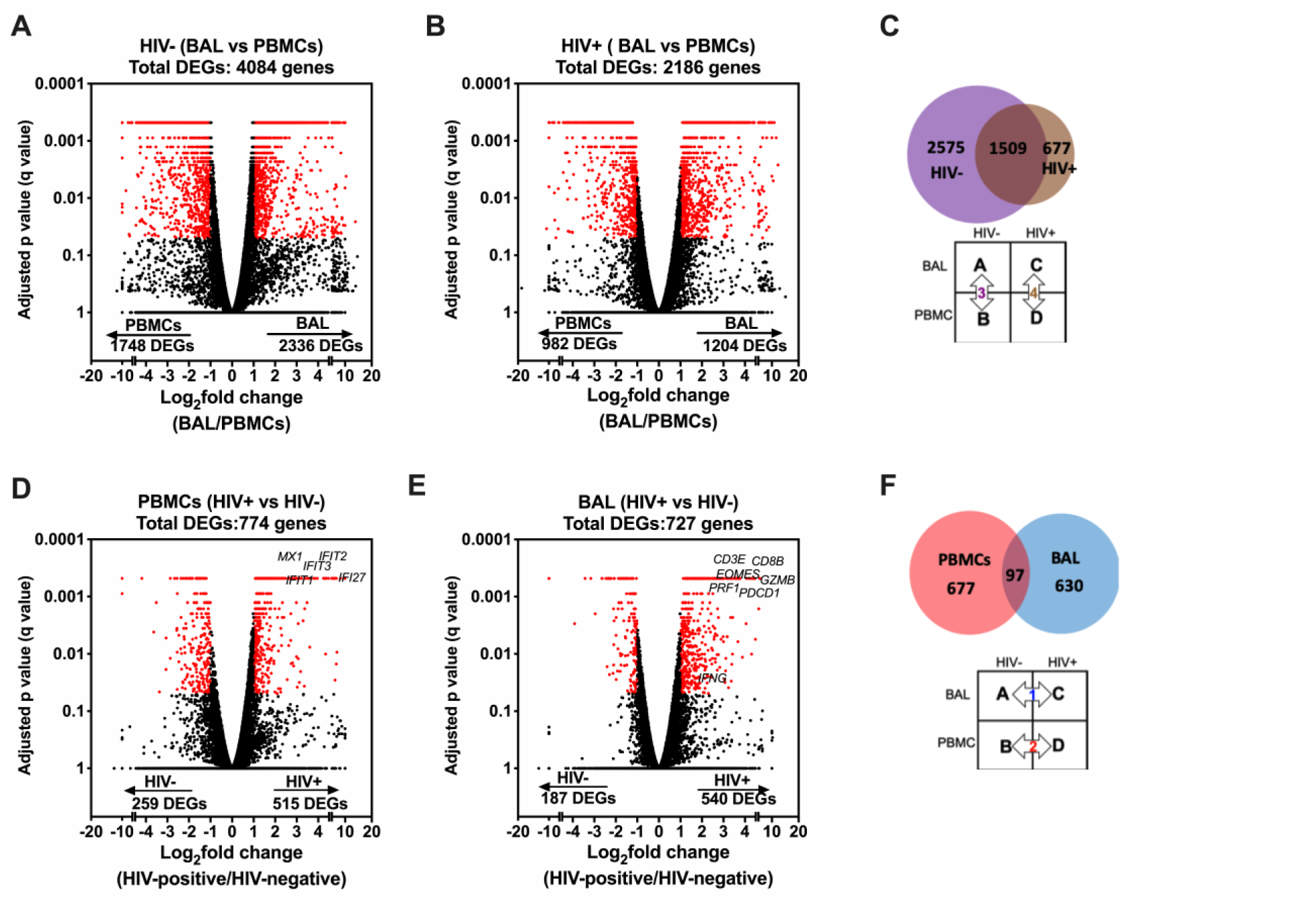
Comparisons in transcriptomic profiles between bronchoalveolar lavage fluid cells (BLCs) and peripheral blood mononuclear cells (PBMCs) in HIV-infected and HIV-uninfected participants. A) Differentially expressed genes between BLCs and PBMCs in the HIV-negative group (n=4). B) Differentially expressed genes between BLCs and PBMCs in the HIV-positive group (n=3). C) Numbers of differentially expressed genes between BLCs and PBMCs in both HIV-positive (n=3) and HIV-negative groups (n=4). D) Differentially expressed genes between HIV-negative (n=4) and HIV-positive (n=3) groups in PBMCs. E) Differentially expressed genes between HIV-negative and HIV-positive groups in BLCs. F) Numbers of differentially expressed genes between HIV-negative and HIV-positive groups in both PBMCs and BLCs.

Using gene ontology (GO) analyses to annotate enriched functions in these compartment specific HIV-induced transcriptional changes, we identified 30 GO terms that were significantly enriched (FDR q-value<0.05) in our list of DEGs in PBMCs between the HIV-positive and HIV-negative groups (Supplementary table 2). We also identified 140 significantly enriched GO terms in BLCs between the HIV-positive and HIV-negative groups (Supplementary table 3). The top enriched GO term in the PBMCs of HIV-positive participants was the “type I interferon signaling pathway” gene set (Figure 3A). On the other hand, the most enriched GO term in the BLCs of HIV-positive participants was the “adaptive immune response” gene set (Figure 3B). We also observed some enrichment of the “response to interferon-beta” and “type I interferon signaling pathway” GO terms in the BLCs of HIV-positive participants, but those GO terms ranked low in our list (Supplementary table 3). Thus, even though HIV also induces a type I interferon signaling signature in bronchoalveolar compartment, its dominant effect there is the modulation of adaptive immune responses.

**Figure 3:**
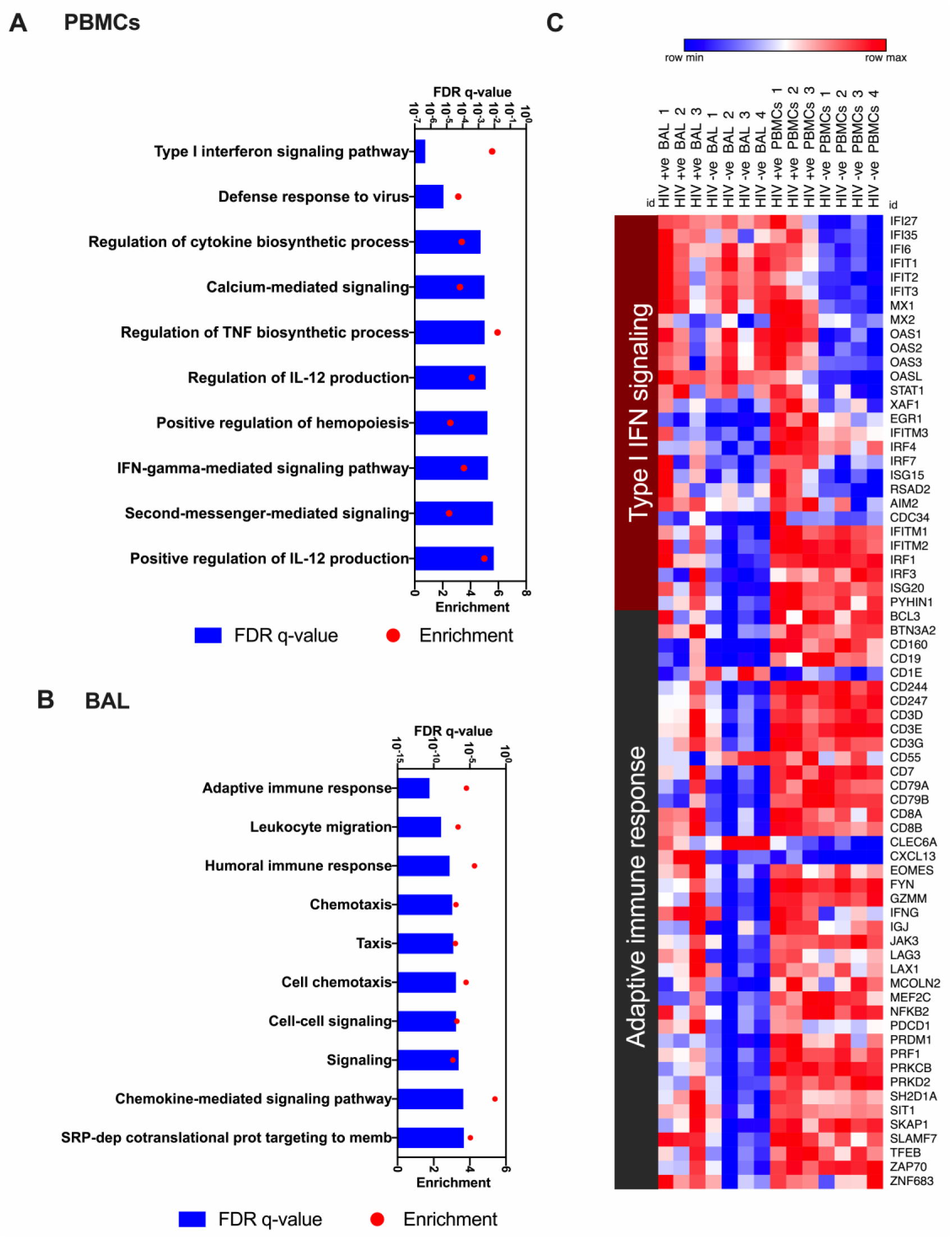
Gene ontology analyses to identify HIV-induced enrichments in bronchoalveolar lavage fluid cells (BLCs) and peripheral blood mononuclear cells (PBMCs). A) The top ten HIV-associated significantly enriched gene ontology (GO) terms in PBMCs. B) The top 10 HIV-associated significantly enriched gene ontology (GO) terms in BLCs. C) Heat map showing expression levels of the genes that contribute to the most enriched GO terms in PBMCs and BLCs i.e. “Type I interferon signaling pathway” and “Adaptive immune response”. Natural logarithms of (Expression level +1) were used.

Enrichment of the “type I interferon signaling pathway” gene set in HIV-positive PBMCs was due to upregulation of twenty genes, namely *EGR1, IFI27, IFI35, IFI6, IFIT1, IFIT2, IFIT3, IFITM3, IRF4, IRF7, ISG15, MX1, MX2, OAS1, OAS2, OAS3, OASL, STAT1, XAF1* and *RSAD2*. On the other hand, enrichments of the “type I interferon signaling pathway” and the “response to interferon-beta” gene sets in HIV-positive BLCs were due to upregulation of fourteen genes, namely *EGR1, IFITM1, IFITM2, IFITM3, IRF1, IRF3, IRF4, IRF7, ISG15, ISG20, RSAD2, AIM2, CDC34* and *PYHIN1* (Figure 3C). Notably, from the above lists of twenty-eight type I interferon-associated genes that were differentially expressed in either of the two compartments, only six (namely *EGR1, IFITM3, IRF4, IRF7, ISG15, and RSAD2*) were differentially expressed in both compartments, suggesting qualitative differences between the compartments.

In BLCs, enrichment of the “adaptive immune response” gene set was attributable to the HIV-induced upregulation of transcripts that suggested infiltration with cytolytic T cells, such as the lineage transcripts for CD8^+^ T cells (*CD3D, CD3E, CD3G, CD8A* and *CD8B)*, CD8^+^ T-cell effector molecules (granzyme M (*GZMM*), perforin (*PRF1*) and interferon gamma (*IFNG*)) and the CD8^+^ T-cell transcription factors (e.g. *EOMES*) (Figure 3C). We also observed contribution by inhibitory/exhaustion markers (PD-1 (*PDCD1*) and LAG-3 (*LAG3*)) in the enrichment of this GO term. Further observation of B-cell transcripts (*CD19, CD79A* and *CD79B*) suggests that HIV also induces infiltration of the bronchoalveolar compartment by other lymphocytes as previously reported ^15^ (Figure 3C).

Since the GO analyses suggested lymphocyte infiltration into the bronchoalveolar compartment, we conducted further targeted analyses on the major lymphocyte lineage markers to determine the infiltrating populations. There were significant upregulations of *CD3D, CD3E, CD3G* and *CD19* transcripts in the BLCs but not PBMCs of HIV-positive patients, suggesting selective HIV-induced expansion in the bronchoalveolar compartment (Supplementary 3A-D). We observed significant upregulations of transcripts for *CD8A* and *CD8B* in both BLCs and PBMCs of HIV-positive patients (Supplementary 3E-F). Even though transcripts for *NCAM1/CD56* were reduced in PBMCs of HIV-positive patients, they were not significantly altered in BLCs, suggesting that the upregulation of cytotoxic markers in the BLCs of HIV-positive patients could be attributed to infiltration by cytotoxic CD8^+^ T cells and not natural killer cells (Supplementary 3G). The levels of *CD4* transcripts did not differ between disease states in either of the compartments, probably because the molecule is expressed on CD4^+^ T cells as well as other lineages such as monocytes and macrophages, a phenomenon that could mask the known HIV-induced CD4^+^ T-cell depletion (Supplementary 3H), with the latter confirmed by flow cytometry in PBMCs (Figure 1E).

The whole compartment transcriptomic approach limited our ability to confidently attribute the transcripts of effector molecules to any specific cell type, and in particular the infiltrating CD8^+^ T cell population which appeared the most likely candidate based on transcriptome profile. We therefore assessed the expression of these molecules in CD8^+^ T cells using flow cytometry and RNA-seq on sorted CD8^+^ T cells. At the protein level, HIV-positive people had higher levels of constitutive Granzyme B production in both PBMCs-derived and BLCs-derived CD8^+^ T cells (Figure 4A). However, *ex vivo* stimulation with mitogens yielded similar inducible interferon gamma between the disease states in both PBMCs-derived and BLCs-derived CD8^+^ T cells (Figure 4B). In agreement with our analyses on bulk BLCs, the HIV-positive individuals had increased expression of PD-1 in BLCs-derived CD8^+^ T cells when compared with HIV-negative individuals (Figure 4C). Similar trends on PD-1 expression were seen when we analyzed the two cohorts separately, confirming that they were immunologically similar (Supplementary figure 4 C and F). Transcriptional analysis of mRNA levels from sorted CD8^+^ T cells confirmed these findings. HIV was associated with a trend of increased transcription for granzyme B (*GZMB)* in BLC-derived CD8^+^ T cells (Figure 4D). Unlike the observation on inducible interferon gamma protein, we observed trends of higher constitutive expression of *IFNG* in HIV-positive individuals in both PBMCs-derived and BLCs-derived CD8^+^ T cells, suggesting that HIV infection could be associated with increased constitutive transcription of *IFNG in vivo* (Figure 4E). In agreement with the data on expression of PD-1 protein on CD8^+^ T cells, we observed a trend of HIV-associated increase in transcription of the PD-1 (*PDCD1*) in both BLCs-derived and PBMCs-derived CD8^+^ T cells (Figure 4F). Thus, the infiltrating CD8+ T cells in HIV-positive individuals’ bronchoalveolar compartment had a functional phenotype consistent with pre-existing cytotoxic products. Considering that PD-1 can be a marker of both exhaustion and activation depending on context, the significance of increased PD-1 expression in BLC-derived CD8 T cells in the HIV-positive group needs to be determined in future studies.

**Figure 4:**
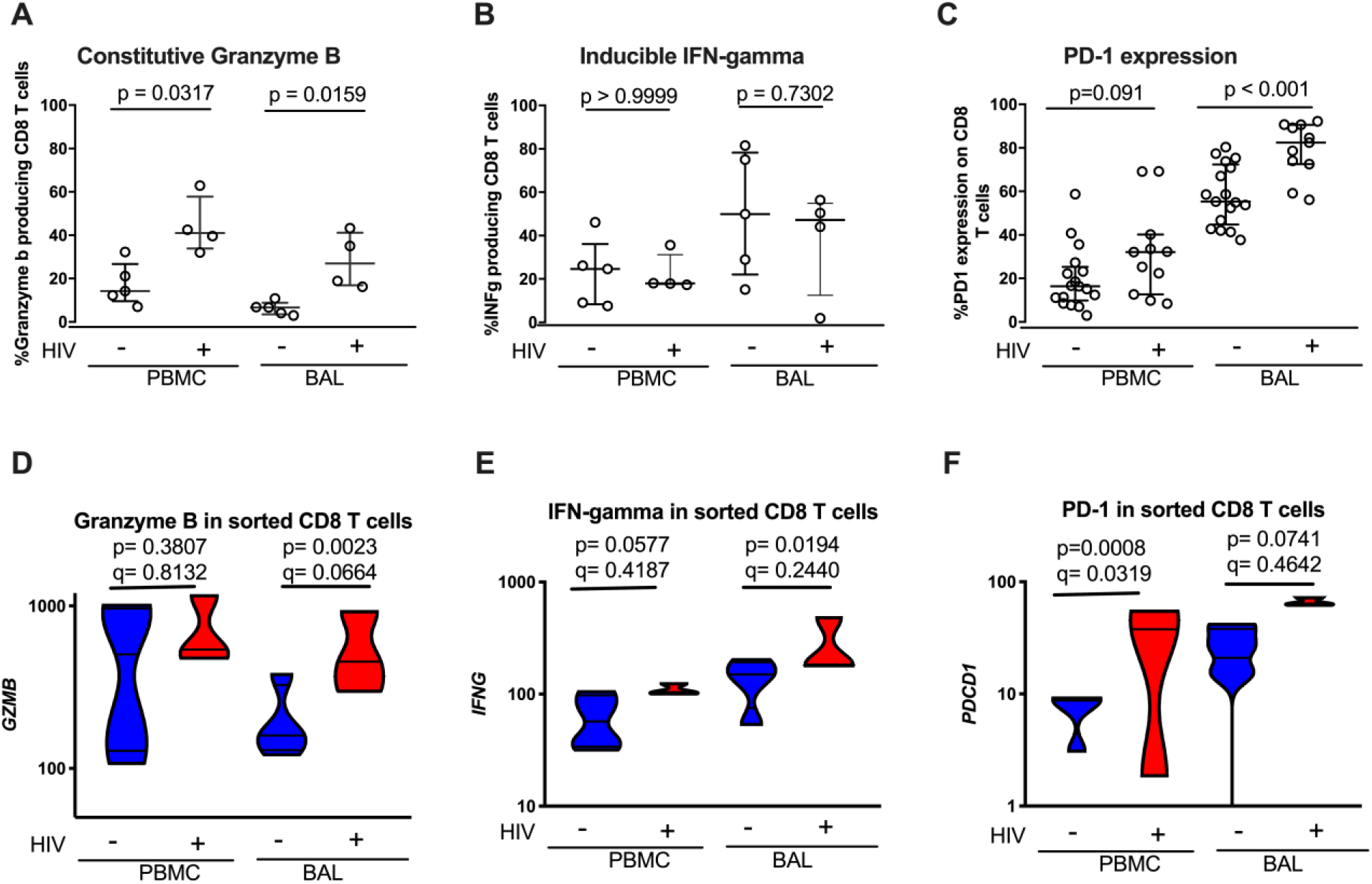
Characterization of HIV-induced infiltrating CD8+ T cells. A) Constitutive expression of granzyme B in unstimulated CD8+ T cells in PBMCs and BLCs from HIV-positive and HIV-negative participants. B) Expression of interferon gamma in unstimulated and stimulated CD8+ T cells in PBMCs and BLCs from HIV-positive and HIV-negative participants. C) *Ex vivo* expression of PD-1 in CD8+ T cells in PBMCs and BLCs from HIV-positive and HIV-negative participants. D) Expression levels of granzyme B mRNA (*GZMB*) in BLCs-derived sorted CD8+ T cells and PBMCs-derived sorted CD8+ T cells from HIV-positive and HIV-negative participants. E) Constitutive expression levels of interferon gamma mRNA (*IFNG*) in BLCs-derived sorted CD8+ T cells and PBMCs-derived sorted CD8+ T cells from HIV-positive and HIV-negative participants. F) Expression levels of PD-1 mRNA (*PDCD1*) in BLCs-derived and PBMCs-derived sorted CD8+ T cells from HIV-positive and HIV-negative participants.

## Discussion

HIV is associated with an increased incidence of both infectious and non-infectious lung morbidities ^13, 16, 17^. HIV-positive patients have a higher prevalence of *Pneumocystis* pneumonia, active tuberculosis, bacterial pneumonia and viral pneumonia ^13, 16^. They also have a higher prevalence of noninfectious structural lung complications such as emphysema and chronic obstructive pulmonary disease (COPD) ^13, 17^.

Despite the high prevalence of HIV-induced lung complications, the immunopathogenesis of HIV in the lung is poorly understood. Due to logistical difficulties of obtaining lung samples, most studies on immune responses to respiratory infections have been conducted in peripheral blood with an assumption that circulating cells have similarities with those in the lung. Here, we show that there are significant differences in the global transcriptional profiles between the blood and the bronchoalveolar compartment, and that the immunological effects of HIV infection revealed by whole compartment transcriptomics in the two compartments are different.

While a type I interferon signature was the most dominant effect of HIV in PBMCs, a CD8^+^ T-cell infiltrate was the dominant effect in the bronchoalveolar compartment. We also observed a weaker and qualitatively different HIV-associated type I interferon signature in the bronchoalveolar compartment of viremic HIV patients. An elevated interferon signature in blood has been associated with progression to TB disease and could arguably relate to the increased susceptibility to TB disease among HIV patients who have a strong type I interferon signature in blood ^10^. The qualitative differences in type I interferon signatures between the compartments could be due to the differences in cellular compositions. Whether specific lung signatures drive the association of type I interferon signaling with progression from latent to active TB remains largely unknown and can be addressed through the study of the bronchoalveolar compartment in participants who go on to develop TB disease.

The infiltration of the bronchoalveolar compartment of HIV-positive patients with CD8^+^ T cells has been reported in previous studies. However, there is limited information on its functional nature ^15, 18^. Using both flow cytometry and whole compartment and CD8^+^ T cell specific RNA-seq, we show that the HIV-induced CD8^+^ T-cell infiltrate is associated with higher expression of effector molecules such as granzymes, perforin and interferon gamma, suggesting a cytolytic and inflammatory profile. We also report higher expression of PD-1 on bronchoalveolar CD8^+^ T cells of HIV-positive patients. PD-1 expression in CD8^+^ T cells has been associated with exhaustion ^19^. However, in juvenile idiopathic arthritis, PD-1^+^ CD8^+^ T cells derived from synovial fluid were metabolically active functional effector memory T cells, suggesting that PD-1 expression could also act as a marker of locally adapted functional T cells ^20^. Thus, depending on context, PD-1 could be a marker of either exhaustion or local activation in tissues. Even though PD-1 blockade on BLCs-derived T cells from HIV-positive patients was previously shown to boost cytokine secretion *in vitro*, suggesting exhaustion, there was a counterintuitive increased PD-1 expression among the interferon gamma secreting T cells ^18^. As such, the implication of HIV-associated increase in PD-1 expression on CD8^+^ T cells in the bronchoalveolar compartment needs further investigation.

The HIV-induced infiltration of the bronchoalveolar compartment with cytolytic CD8^+^ T cells could be driven directly by HIV replication ^21^. The lung has been shown to be a site of HIV replication where small alveolar macrophages and CCR5 expressing CD4^+^ T cells are preferentially infected with HIV ^9, 22^. Infiltrating CD8^+^ T cells could control local HIV replication by killing the HIV-infected CD4^+^ T cells and alveolar macrophages. Indeed, in previous studies, lymphocytic alveolitis in asymptomatic HIV patients was enriched for HIV-specific cytotoxic CD8^+^ T cells that could execute such effector functions ^18^. Whether HIV induces bronchoalveolar infiltration with other specificities of CD8^+^ T cells that can modulate opportunistic respiratory infections, such as tuberculosis, is unclear.

In other settings, CD8 lymphocytic alveolitis has been implicated in the pathogenesis of noninfectious lung complications, such as COPD and emphysema. Considering that HIV infection is also associated with increased prevalence of the same noninfectious lung complications, HIV-induced lymphocytic alveolitis is thought to accelerate the deterioration of lung function in patients who are exposed to other risk factors for COPD and emphysema, such as smokers ^21, 23-26^. Whether similar bystander destructive mechanisms play an important role in CD8^+^ T-cell-mediated disruption of the containment of *Mycobacterium tuberculosis* in granulomas is unknown. In an immune-competent mouse model, LCMV-specific CD8^+^ T cells infiltrated *Mycobacterium bovis* granulomas in the liver, but without conferring any benefit in the control of bacterial growth, suggesting that HIV-specific CD8^+^ T cells in our setting could also infiltrate *Mycobacterium tuberculosis* granulomas in human hosts ^27^. In another mouse model, LCMC-specific cytolytic CD8^+^ T cells expressing granzyme B and NKG2D infiltrated *Leishmania major* lesions and exacerbated disease by causing an exaggerated inflammatory response ^28^. Infiltrating cytolytic CD8^+^ T cells in the lung of HIV patients could similarly exaggerate the inflammatory state, disrupting the containment of *Mycobacterium tuberculosis* in granulomas thus promoting bacterial dissemination. Notably, the CD8^+^ T-cell infiltrate in our cohorts was characterized by increased expression of granzymes and perforin, suggesting some overlap between the findings in our cohorts and the *Leishmania major* mouse model ^28^. Additional studies will be needed to directly interrogate the possible contribution of infiltrating CD8^+^ T cells in the inflammatory destruction of lung tissues and the anatomical dissemination of *Mycobacterium tuberculosis* infections.

We conclude that HIV is associated with a cytolytic CD8^+^ T-cell infiltrate in the bronchoalveolar compartment. Further mechanistic studies are required to understand the consequences of the infiltration on respiratory infections, such as *Mycobacterium tuberculosis*, and noninfectious comorbidities, such as COPD and emphysema. Our study did not assess the antigen specificity of the infiltrating CD8^+^ T cells, although a previous report suggested an enrichment for HIV-specific CD8^+^ T cells ^18^. In future studies, it will be important to determine whether enrichment for CD8^+^ T cells against respiratory infections, such as *Mycobacterium tuberculosis* occurs, and the functional competence of these cell. This study was limited by the sample size in the transcriptomic profiling and intracellular cytokine staining. Nevertheless, the data reveal important compartment-specific effects of HIV in the bronchoalveolar compartment, suggesting a possible mechanism by which HIV modulates immunity to respiratory infections and lung function in ways that cannot be revealed by studying peripheral blood. Furthermore, we show the utility of using whole compartment transcriptomic analyses to reveal infiltration of different sites with various immune cells.

## Methods

### Study population

We studied the effect of HIV on immune function in the peripheral blood and bronchoalveolar compartment using two bronchoalveolar study cohorts at African Health Research Institute (AHRI) in KwaZulu-Natal, South Africa. The first cohort was a hospital-based cohort in which we recruited HIV-negative or HIV-positive ART-naive participants (>18yrs) who came for clinical investigations but were determined (after extensive work-up including bronchoscopy and bronchoalveolar microbiological investigations) to not have any infectious or inflammatory pulmonary disease. Their clinical indications and final diagnosis are documented in supplemental table 4. Participants were consented for research use of clinically excess BAL fluid and a paired peripheral blood draw. The second cohort was a research bronchoscopy cohort of HIV-negative and HIV-positive adults (18-50yrs). Exclusion criteria included: pregnancy, any history of disease other than HIV, history of antiretroviral therapy (ART), and smoking. Study participants were recruited from KwaDabeka Community Health Centre. All participants were confirmed to be free of respiratory symptoms and to have a normal chest x-ray. Further, HIV-positive participants were confirmed to have a negative sputum Mtb GeneXpert. The HIV status of all participants was determined by 4th generation HIV antibody/antigen Enzyme Linked-Immunosorbent Assay (ELISA) testing and HIV RNA quantitative viral load. CD4^+^ T cell counts were determined in all participants. All participants also underwent assessment of hemoglobin (had at least 10g/dL), platelet level (had at least 100×10^9^ cell/L) and prothrombin time (INR<1.3) to meet safety criteria for bronchoscopy. Once screened and characterized, participants were transported to Inkosi Albert Luthuli Central Hospital (IALCH) where they underwent research bronchoscopy and paired peripheral blood draw. All participants provided written informed consent. Both study protocols were approved by the University of KwaZulu-Natal Biomedical Research Ethics Committee (BREC; reference numbers BF503/15 and BE037/12) and Partners Institutional Review Board.

Participant samples were selectively subjected (depending on sample availability for different techniques) to differential cell count, mitogen stimulation, monoclonal antibody staining, and transcriptomic analysis.

### Sample processing and differential cell counts

Bronchoscopies were performed by pulmonologists at IALCH with participants receiving sedation and bronchodilators according to local standard of care protocols. Two hundred milliliters of normal saline were infused into the right middle lobe. Bronchoalveolar lavage fluid was stored at 4 °C and processed in the laboratory within 90 minutes. Paired peripheral blood was collected in acid citrate dextrose (ACD) tubes (BD, Franklin Lakes, NJ, USA) and stored at room temperature. A portion of the sample was directly used for differential cell counts in each compartment after standard preparation and interpretation of peripheral blood smear and cytospin slide for BAL fluid. PBMCs were isolated using standard Histopaque (Sigma-Aldrich, St. Louis, MO, USA) gradient centrifugation protocols.

To isolate bronchoalveolar lavage fluid cells (BLCs), BAL fluid was passed through a 40 µm filter (BD). The fluid was spun at 1,500 RPM for 10 minutes at 4 °C and all cells resuspended in RPMI media supplemented with 5% fetal bovine serum, 1% penicillin/streptavidin, 1% HEPES buffer and 1% amphotericin. The cells were freshly used for monoclonal antibody staining and mitogen stimulation to test functionality.

### Flow cytometry

The BLCs and PBMCs were counted and assessed for viability using trypan blue (Sigma-Aldrich) and compound microscopy to ensure > 90% lymphocyte viability. To assess distribution of CD4^+^ and CD8^+^ T cells, immune regulation and functionality of CD8^+^ T cells in HIV infection, cells from the two compartments were subjected to two panels of fluorescently labelled antibodies. Panel 1 (phenotypic panel): Live/dead-amcyan (Life technologies, Carlsbad, CA, USA), CD3-BV650, CD4-BV711, CD14-APC-Cy7, PD-1-BV711, TIM-3-BV785 (Biolegend, San Diego, CA, USA) and CD8-PE Texas Red (Invitrogen, Carlsbad, CA, USA). Panel 2 (intracellular cytokine staining panel): Live/dead-amcyan (Life technologies), CD3-Alexa700, CD4-BV711, CD8-APC-Cy7, granzyme b-Alexa647, IFNγ-Dazzle 549 (Biolegend). Panel 2 staining was done after stimulation with PMA/Ionomycin (25/500ng/mL). Acquisition was performed on BD FACS Aria III. Flowjo v10.5 (Flowjo, LLC) was used for flow cytometry analysis.

### RNA isolation from Whole BAL

Freshly processed cellular pellets from bronchoalveolar lavage (1 mL of BAL fluid) and Histopaque gradient-isolated PBMCs (1 × 10^6^ cells) from the hospital cohort for bulk sequencing were stored in RNAlater stabilizing reagent (Sigma-Aldrich) at −80 °C. Samples were later thawed at room temperature, pelleted and suspended with 1% β-mercaptoethanol RLT buffer from the Qiagen RNeasy Micro kit (Qiagen, Hilden, Germany). Extraction of RNA was performed according to manufacturer’s protocol. Briefly, a QIAshredder column was used to homogenize the samples. DNAse 1 treatment was used to eliminate any remaining genomic DNA contamination. The extracted RNA was quantified using nanodrop and aliquoted into ∼200 ng aliquots, adequate for RNASeq library preparation. All aliquots were immediately stored at −80 °C.

### Sorted CD8+ T cell populations

For work on purified CD8^+^ T cells, freshly processed cells from the hospital cohort were sorted into 70% TRIzol LS Reagent (Thermo Fisher Scientific, Waltham, MA, USA) and stored at −80 °C. Both RNA and DNA were isolated from these samples using the TRIzol/chloroform method with minor modifications. Briefly, RNA was precipitated using a nucleic acid co-precipitant, 5 mg/mL linear acrylamide (Thermo Fisher Scientific). All reagents were kept at 4 °C to facilitate separation of nucleic acid into different layers and efficient precipitation. QIAGEN RNeasy Micro kit was used to purify RNA from the TRIzol extracted samples. The quality of the extracted RNA was determined using the BioAnalyzer RNA Pico kit (Agilent, Santa Clara, CA, USA).

### RNA-seq library preparation and sequencing

Enrichment for messenger RNA was done using the NEBNext Poly(A) mRNA Magnetic Isolation kit (New England Biolabs, Ipswich, MA, USA). RNA libraries were prepared using the NEBNext Ultra RNA Library Prep Kit for Illumina (New England Biolabs). Dual index primers from the NEBNext Multiplex Oligos for Illumina kit were used to label the samples. A subset of the libraries was assessed for acceptable quality using the BioAnalyzer DNA High Sensitivity Chip or the DNA TapeStation (Agilent). Concentrations of the libraries were determined using a Qubit dsDNA assay kit (Thermo Fisher). Equal molarities of the indexed libraries were pooled and sequenced on an Illumina NextSeq 500 platform to yield 75 bp paired end reads.

### Sequencing data analyses

The raw data were demultiplexed and processed using Trimmomatic version 0.36 to remove adaptors and leading/trailing low-quality bases. Subsequent analyses were done using the Tuxedo protocol as previously described ^29^. Briefly, the sequences were aligned on the human reference genome *GRCh37* (hg19) using the TopHat module (version 2.1.1) and bowtie (version 2.2.4). The mapped reads were then sorted using the Picard SortSam module. Duplicate reads were identified using the Picard MarkDuplicates module and removed. The Cufflinks module (Version 2.2.1) was used for subsequent analyses. The transcripts for each sample were assembled and the numbers of reads quantified for each transcript. The expression levels were expressed as Fragments Per Kilobase of transcript per Million fragments mapped (FPKM). The assembled transcripts for all samples were merged to obtain a master transcriptome assembly (Cuffmerge) followed by assessment of differential expression (Cuffdiff). A statistically significant difference in the expression of a transcript between two groups of participants was defined as having at least a two-fold difference and q < 0.05 (after Benjamini-Hochberg correction for multiple-testing). CummeRbund R package (version 2.16), GraphPad Prism (version 8) and Morpheus-Broad Institute (https://software.broadinstitute.org/morpheus/) were used for subsequent data visualization. Identification of HIV-induced pathway changes was done on the GOrilla platform http://cbl-gorilla.cs.technion.ac.il/) by checking for enriched Gene Ontology (GO) terms among the differentially expressed genes ^30^. To exclude very large or very small nonspecific GO terms that did not have specific biological implications, an arbitrary cut-off was set whereby only GO terms whose sizes are between 20 and 200 genes were considered in the analyses.

### Statistical analyses

Comparisons of flow cytometry data and differential counts data between HIV-uninfected and HIV-infected groups were assessed using the Wilcoxon rank-sum test. Matched comparisons of flow cytometry data and differential counts data between blood and bronchoalveolar lavage samples were assessed using the Wilcoxon matched pairs signed rank test. Differences were considered statistically significant if p < 0.05. Comparisons of ratios of participants between groups were done using Chi-square test. All statistical analyses were done on GraphPad Prism version 8 (GraphPad Software, Inc).

## Acknowledgements

This work was supported in part by grants from the South African Medical Research Council, the Department of Science and Technology/National Research Foundation Research Chairs Initiative, the Victor Daitz Foundation, a Burroughs-Wellcome Fund / American Society of Tropical Medicine and Hygiene fellowship, the National Institutes of Health (K08 AI118538), the Harvard University Center for AIDS Research (grant P30 AI060354), a capacity building Strategic Award awarded to the KEMRI-Wellcome Trust Research Programme and the South Africa National Research Foundation. Additional support was received through the Saharan African Network for TB/HIV Research Excellence (SANTHE), a DELTAS Africa Initiative [grant # DEL-15-006]. The DELTAS Africa Initiative is an independent funding scheme of the African Academy of Sciences (AAS)’s Alliance for Accelerating Excellence in Science in Africa (AESA) and supported by the New Partnership for Africa’s Development Planning and Coordinating Agency (NEPAD Agency) with funding from the Wellcome Trust [grant # 107752/Z/15/Z] and the UK government. The views expressed in this publication are those of the author(s) and not necessarily those of AAS, NEPAD Agency, Wellcome Trust or the UK government.

The authors declare no conflict of interest.

## Authors contributions

D.M.M., Maphe Mthembu, A.S., U.S., S.S.R., B.C. and T.B. performed the experiments. K.N., D.F.K., P.M., Mohammed Mitha, M.S., Z.M., D.R., F.K. and Tarryn Naidoo collected the clinical samples. D.K., Thumbi Ndung’u and E.B.W. provided supervision. D.M.M., Maphe Mthembu, Thumbi Ndung’u and E.B.W. wrote the manuscript. All authors reviewed the manuscript and approved the final version.

## Supplementary material

All supplementary material and gene expression data are available at Figshare: https://figshare.com/s/e05711feb0360fe3ce33

## Disclosure

The authors declare no competing interests.

## Notes

https://figshare.com/s/e05711feb0360fe3ce33

