## Supplementary figures for "Contrasting inflammatory signatures in peripheral blood and bronchoalveolar cells reveal compartment-specific effects of HIV infection"

### Supplementary material

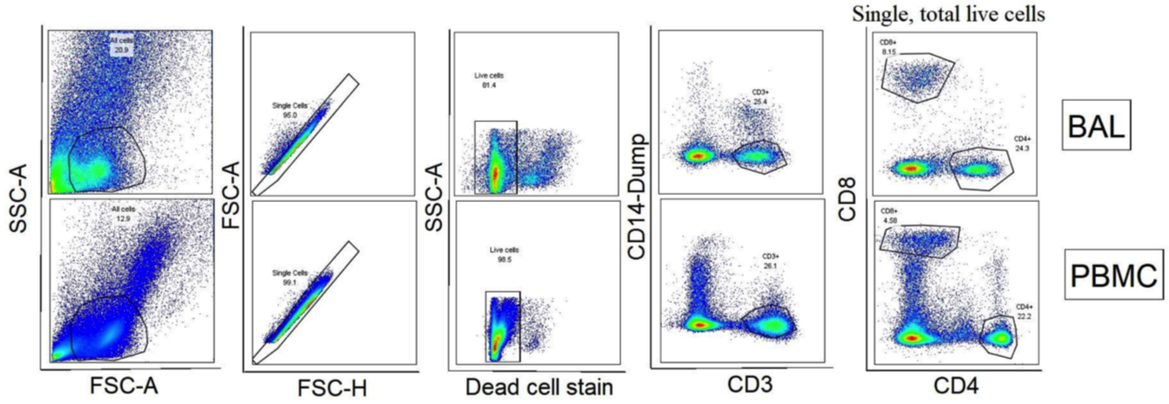


**Supplementary Figure 1: Gating strategy for sorting and characterizing CD8$^+$ T cells.**

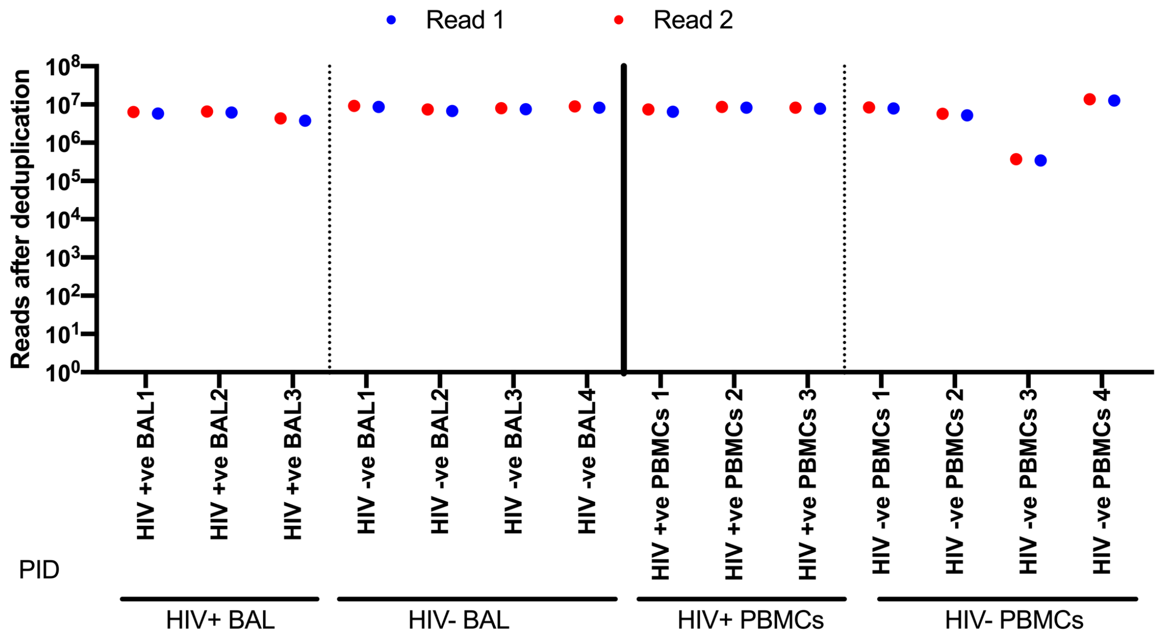


**Supplementary Figure 2: Numbers of unique forward and reverse reads for each sample after deduplication.**

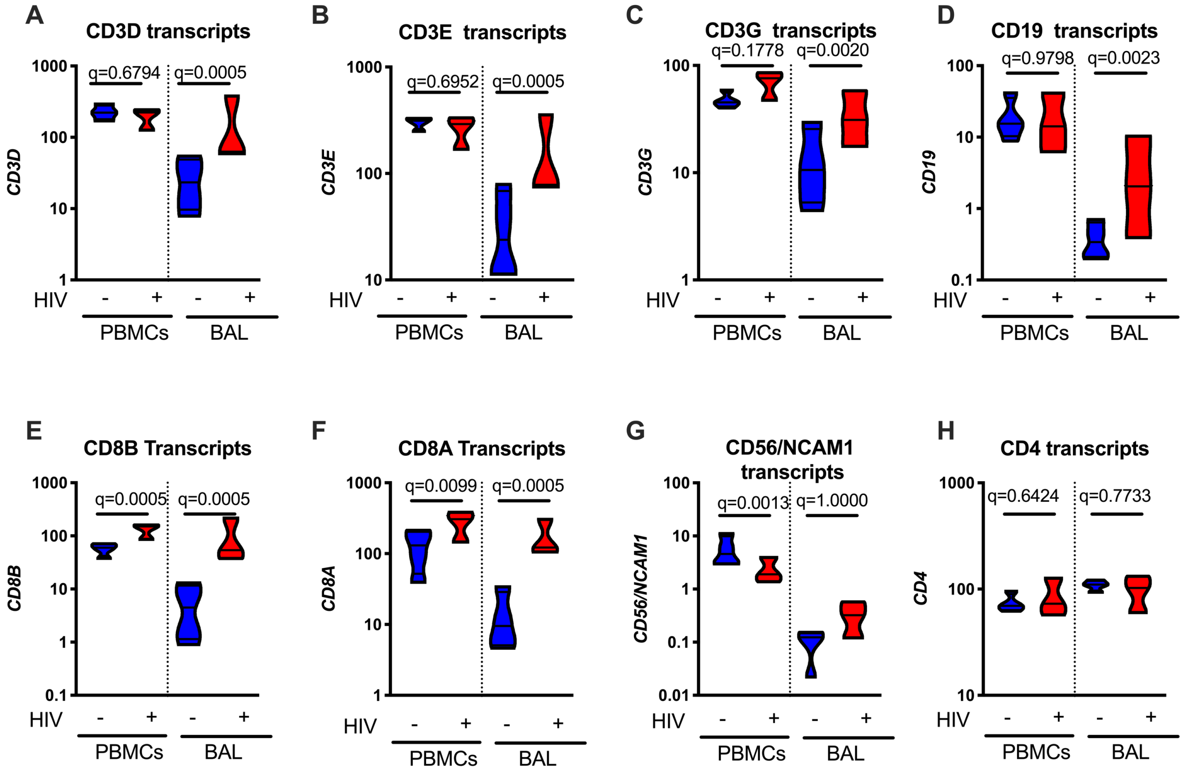


**Supplementary Figure 3: Characterization of the effect of HIV on distribution of lymphocytes lineages in BLCs and PBMCs**. A) Expression of *CD3D* in whole BLCs and PBMCs in HIV-positive and HIV-negative groups. B) Expression of *CD3E* in whole BLCs and PBMCs in HIV-positive and HIV-negative groups. C) Expression of *CD3G* in whole BLCs and PBMCs in HIV-positive and HIV-negative groups. D) Expression of *CD19* in whole BLCs and PBMCs in HIV-positive and HIV-negative groups. E) Expression of *CD8B* in whole BLCs and PBMCs in HIV-positive and HIV-negative groups. F) Expression of *CD8A* in whole BLCs and PBMCs in HIV-positive and HIV-negative groups. G) Expression of *CD56/NCAM1* in whole BLCs and PBMCs in HIV-positive and HIV-negative groups. H) Expression of *CD4* in whole BLCs and PBMCs in HIV-positive and HIV-negative groups.

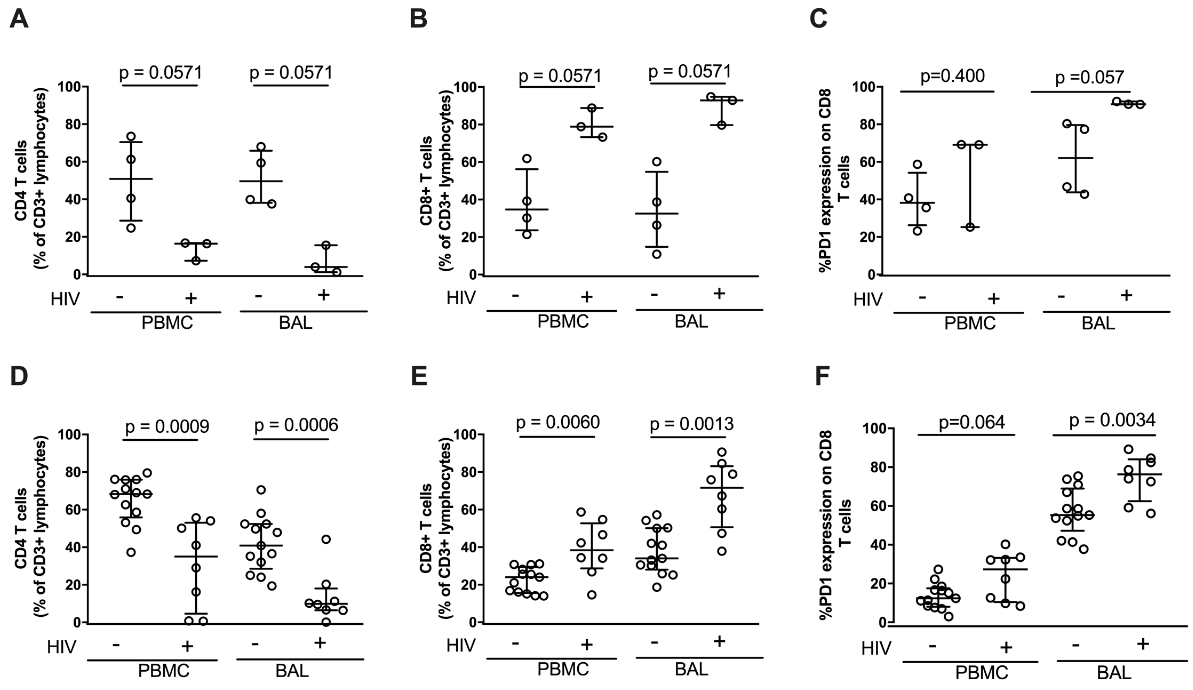


A-C: Hospital-based cohort
D-F: Research bronchoscopy cohort

**Supplementary Figure 4: Comparison of the two bronchoscopy cohorts HIV infection effect in T cell distribution and phenotype.** A-C: Shows the hospital-based bronchoscopy cohort CD4 T cells, CD8 T cells distribution and CD8 T cells PD-1 expression in the peripheral blood and bronchoalveolar compartment in HIV-negative and HIV-positive people. D-F: Shows the Research bronchoscopy cohort CD4 T cells, CD8 T cells distribution and CD8 T cells PD-1 expression in the peripheral blood and bronchoalveolar compartment in HIV-negative and HIV-positive people.
