## Supplementary Tables for "Contrasting inflammatory signatures in peripheral blood and bronchoalveolar cells reveal compartment-specific effects of HIV infection"

**Supplementary Table 1: Immune cell subsets composition between the peripheral blood and bronchoalveolar compartment**

| | Bloods | | | BAL | | | p-value blood vs BAL | |
|---|---|---|---|---|---|---|---|---|
| **Cell types** | **HIV-** | **HIV+** | **p-value** | **HIV-** | **HIV+** | **p-value** | **HIV-** | **HIV+** |
| % Macrophages/monocytes | 6.9 (6.2-9.35) | 8.45 (7.2-9.3) | 0.456 | 87.4 (79.8-90) | 79.25 (70.9-88) | 0.258 | <0.0001 | 0.0078 |
| % Neutrophils | 45.4 (36.8-57) | 50.35 (36.9-65.5) | 0.591 | 3.2 (1.4-5.4) | 1 (0.7-1.7) | 0.183 | <0.0001 | 0.0078 |
| % Eosinophils | 1.6 (1.1-3.05) | 1.95 (1.8-2.8) | 0.418 | 0 (0-1) | 0.3 (0.15-0.85) | 0.413 | 0.0424 | 0.0156 |
| % Lymphocytes | 43.4 (34.15-50.6) | 37.8 (22.2-46.5) | 0.428 | 7.8 (5-14) | 8.8 (7.7-23.9) | 0.364 | <0.0001 | 0.039 |
| Proportion CD3+ T lymphocytes | 17.6 (15.29-23.65) | 5.9 (2.87-13.16) | 0.003 | 0.59 (0.33-2.80) | 3.02 (1.48-6.10) | 0.0471 | 0.0005 | 0.25 |
| Proportion CD4+ T lymphocytes | 68 (53.1-73.5) | 16.7 (13.97-45.55) | 0.0001 | 40.9 (35.2-52.4) | 9.52 (5.17-13.35) | <0.0001 | 0.0131 | 0.0537 |
| Proportion CD8+ T lymphocytes | 25.5 (16.9-30.7) | 46.6 (34.25-66) | 0.002 | 34 (26.4-50) | 78.9 (63.85-87.55) | <0.0001 | 0.0519 | 0.0068 |

Data presented in median and interquartile range unless stated otherwise. Mann Whitney and Wilcoxon tests were used to calculate significance between the two study groups i.e. HIV- and HIV+, and within each compartment, respectively. ns means p value > 0.05.

**Supplementary Table 2: Enriched GO terms in the list of genes that were differentially expressed in PBMCs between the HIV-positive and HIV-negative participants.** Go terms that had 20-200 genes were considered.

| GO Term | Description | P-value | FDR q-value | Enrichment | N | B | n | b |
|---|---|---|---|---|---|---|---|---|
| GO:0060337 | type I interferon signaling pathway | 6.68E-11 | 4.50E-07 | 5.55 | 9802 | 50 | 707 | 20 |
| GO:0051607 | defense response to virus | 4.62E-09 | 6.23E-06 | 3.12 | 9802 | 142 | 707 | 32 |
| GO:0042035 | regulation of cytokine biosynthetic process | 3.52E-06 | 1.35E-03 | 3.37 | 9802 | 74 | 707 | 18 |
| GO:0019722 | calcium-mediated signaling | 6.45E-06 | 2.41E-03 | 3.24 | 9802 | 77 | 707 | 18 |
| GO:0042534 | regulation of tumor necrosis factor biosynthetic process | 6.70E-06 | 2.44E-03 | 5.94 | 9802 | 21 | 707 | 9 |
| GO:0032655 | regulation of interleukin-12 production | 8.19E-06 | 2.83E-03 | 4.1 | 9802 | 44 | 707 | 13 |
| GO:1903708 | positive regulation of hemopoiesis | 1.16E-05 | 3.64E-03 | 2.55 | 9802 | 136 | 707 | 25 |
| GO:0060333 | interferon-gamma-mediated signaling pathway | 1.28E-05 | 3.93E-03 | 3.52 | 9802 | 59 | 707 | 15 |
| GO:0019932 | second-messenger-mediated signaling | 3.08E-05 | 8.14E-03 | 2.46 | 9802 | 135 | 707 | 24 |
| GO:0032735 | positive regulation of interleukin-12 production | 3.58E-05 | 9.11E-03 | 4.99 | 9802 | 25 | 707 | 9 |
| GO:0009887 | animal organ morphogenesis | 3.52E-05 | 9.11E-03 | 2.16 | 9802 | 199 | 707 | 31 |
| GO:0048534 | hematopoietic or lymphoid organ development | 5.57E-05 | 1.32E-02 | 2.55 | 9802 | 114 | 707 | 21 |
| GO:0030198 | extracellular matrix organization | 5.85E-05 | 1.36E-02 | 2.62 | 9802 | 106 | 707 | 20 |
| GO:0045637 | regulation of myeloid cell differentiation | 8.04E-05 | 1.75E-02 | 2.24 | 9802 | 161 | 707 | 26 |
| GO:1902105 | regulation of leukocyte differentiation | 7.94E-05 | 1.75E-02 | 2.1 | 9802 | 198 | 707 | 30 |
| GO:0070542 | response to fatty acid | 8.99E-05 | 1.92E-02 | 3.54 | 9802 | 47 | 707 | 12 |
| GO:0032103 | positive regulation of response to external stimulus | 9.71E-05 | 2.04E-02 | 2.18 | 9802 | 172 | 707 | 27 |

| | | | | | | | | |
|---|---|---|---|---|---|---|---|---|
| GO:0043901 | negative regulation of multi-organism process | 1.24E-04 | 2.49E-02 | 2.36 | 9802 | 129 | 707 | 22 |
| GO:0032647 | regulation of interferon-alpha production | 1.30E-04 | 2.58E-02 | 4.82 | 9802 | 23 | 707 | 8 |
| GO:0001505 | regulation of neurotransmitter levels | 1.57E-04 | 2.94E-02 | 2.28 | 9802 | 140 | 707 | 23 |
| GO:0032652 | regulation of interleukin-1 production | 1.61E-04 | 2.97E-02 | 3.16 | 9802 | 57 | 707 | 13 |
| GO:0032651 | regulation of interleukin-1 beta production | 2.09E-04 | 3.81E-02 | 3.26 | 9802 | 51 | 707 | 12 |
| GO:0042108 | positive regulation of cytokine biosynthetic process | 2.54E-04 | 4.13E-02 | 3.2 | 9802 | 52 | 707 | 12 |
| GO:0001101 | response to acid chemical | 2.54E-04 | 4.17E-02 | 2.06 | 9802 | 182 | 707 | 27 |
| GO:0032680 | regulation of tumor necrosis factor production | 2.64E-04 | 4.23E-02 | 2.42 | 9802 | 109 | 707 | 19 |
| GO:0030097 | hemopoiesis | 2.78E-04 | 4.36E-02 | 3 | 9802 | 60 | 707 | 13 |
| GO:1903428 | positive regulation of reactive oxygen species biosynthetic process | 3.11E-04 | 4.56E-02 | 3.9 | 9802 | 32 | 707 | 9 |
| GO:0042116 | macrophage activation | 3.11E-04 | 4.61E-02 | 3.9 | 9802 | 32 | 707 | 9 |
| GO:1903555 | regulation of tumor necrosis factor superfamily cytokine production | 3.35E-04 | 4.71E-02 | 2.37 | 9802 | 111 | 707 | 19 |
| GO:0045428 | regulation of nitric oxide biosynthetic process | 3.33E-04 | 4.72E-02 | 3.55 | 9802 | 39 | 707 | 10 |

**Supplementary Table 3: Enriched GO terms in the list of genes that were differentially expressed in BLCs between the HIV-positive and HIV-negative participants.** Go terms that had 20-200 genes were considered.

| GO Term | Description | P-value | FDR q-value | Enrichment | N | B | n | b |
|---|---|---|---|---|---|---|---|---|
| GO:0002250 | adaptive immune response | 1.53E-14 | 2.59E-11 | 3.81 | 9950 | 164 | 668 | 42 |
| GO:0050900 | leukocyte migration | 9.29E-13 | 1.05E-09 | 3.35 | 9950 | 191 | 668 | 43 |
| GO:0006959 | humoral immune response | 2.39E-11 | 1.61E-08 | 4.26 | 9950 | 98 | 668 | 28 |
| GO:0006935 | chemotaxis | 6.38E-11 | 3.75E-08 | 3.23 | 9950 | 175 | 668 | 38 |
| GO:0042330 | taxis | 9.12E-11 | 5.13E-08 | 3.2 | 9950 | 177 | 668 | 38 |
| GO:0060326 | cell chemotaxis | 2.32E-10 | 1.21E-07 | 3.79 | 9950 | 114 | 668 | 29 |
| GO:0007267 | cell-cell signaling | 2.48E-10 | 1.24E-07 | 3.28 | 9950 | 159 | 668 | 35 |
| GO:0023052 | signaling | 5.99E-10 | 2.79E-07 | 3.06 | 9950 | 180 | 668 | 37 |
| GO:0070098 | chemokine-mediated signaling pathway | 3.82E-09 | 1.26E-06 | 5.39 | 9950 | 47 | 668 | 17 |
| GO:0006614 | SRP-dependent cotranslational protein targeting to membrane | 4.73E-09 | 1.49E-06 | 4.03 | 9950 | 85 | 668 | 23 |
| GO:0030595 | leukocyte chemotaxis | 7.72E-09 | 2.32E-06 | 3.94 | 9950 | 87 | 668 | 23 |
| GO:0030098 | lymphocyte differentiation | 1.12E-08 | 3.28E-06 | 3.1 | 9950 | 149 | 668 | 31 |
| GO:0006613 | cotranslational protein targeting to membrane | 1.56E-08 | 4.40E-06 | 3.81 | 9950 | 90 | 668 | 23 |
| GO:0000184 | nuclear-transcribed mRNA catabolic process, nonsense-mediated decay | 1.95E-08 | 5.38E-06 | 3.43 | 9950 | 113 | 668 | 26 |
| GO:0009617 | response to bacterium | 6.80E-08 | 1.70E-05 | 2.94 | 9950 | 152 | 668 | 30 |
| GO:0045047 | protein targeting to ER | 7.12E-08 | 1.75E-05 | 3.53 | 9950 | 97 | 668 | 23 |
| GO:0050921 | positive regulation of chemotaxis | 7.96E-08 | 1.86E-05 | 3.77 | 9950 | 83 | 668 | 21 |
| GO:0072599 | establishment of protein localization to endoplasmic reticulum | 1.30E-07 | 2.92E-05 | 3.43 | 9950 | 100 | 668 | 23 |
| GO:0072676 | lymphocyte migration | 2.13E-07 | 4.63E-05 | 5.09 | 9950 | 41 | 668 | 14 |

| | | | | | | | | |
|---|---|---|---|---|---|---|---|---|
| GO:0006968 | cellular defense response | 2.74E-07 | 5.69E-05 | 5.38 | 9950 | 36 | 668 | 13 |
| GO:0050920 | regulation of chemotaxis | 3.40E-07 | 6.86E-05 | 3.08 | 9950 | 121 | 668 | 25 |
| GO:1990266 | neutrophil migration | 3.59E-07 | 7.13E-05 | 4.33 | 9950 | 55 | 668 | 16 |
| GO:0030593 | neutrophil chemotaxis | 3.99E-07 | 7.82E-05 | 4.56 | 9950 | 49 | 668 | 15 |
| GO:0019083 | viral transcription | 4.77E-07 | 8.84E-05 | 3.2 | 9950 | 107 | 668 | 23 |
| GO:0007204 | positive regulation of cytosolic calcium ion concentration | 7.61E-07 | 1.35E-04 | 2.96 | 9950 | 126 | 668 | 25 |
| GO:0070972 | protein localization to endoplasmic reticulum | 8.02E-07 | 1.37E-04 | 3.11 | 9950 | 110 | 668 | 23 |
| GO:0097530 | granulocyte migration | 7.97E-07 | 1.38E-04 | 4.11 | 9950 | 58 | 668 | 16 |
| GO:0071621 | granulocyte chemotaxis | 9.40E-07 | 1.46E-04 | 4.3 | 9950 | 52 | 668 | 15 |
| GO:0032103 | positive regulation of response to external stimulus | 9.76E-07 | 1.50E-04 | 2.52 | 9950 | 189 | 668 | 32 |
| GO:0045071 | negative regulation of viral genome replication | 1.06E-06 | 1.59E-04 | 4.53 | 9950 | 46 | 668 | 14 |
| GO:0002687 | positive regulation of leukocyte migration | 2.10E-06 | 2.90E-04 | 3.37 | 9950 | 84 | 668 | 19 |
| GO:0045619 | regulation of lymphocyte differentiation | 2.34E-06 | 3.19E-04 | 2.86 | 9950 | 125 | 668 | 24 |
| GO:0046634 | regulation of alpha-beta T cell activation | 3.86E-06 | 4.88E-04 | 3.52 | 9950 | 72 | 668 | 17 |
| GO:0048525 | negative regulation of viral process | 3.86E-06 | 4.92E-04 | 3.52 | 9950 | 72 | 668 | 17 |
| GO:0042110 | T cell activation | 4.44E-06 | 5.41E-04 | 2.63 | 9950 | 147 | 668 | 26 |
| GO:0009306 | protein secretion | 5.25E-06 | 6.16E-04 | 3.18 | 9950 | 89 | 668 | 19 |
| GO:0002790 | peptide secretion | 5.23E-06 | 6.20E-04 | 3.07 | 9950 | 97 | 668 | 20 |
| GO:0002685 | regulation of leukocyte migration | 5.23E-06 | 6.25E-04 | 2.81 | 9950 | 122 | 668 | 23 |
| GO:0006612 | protein targeting to membrane | 6.31E-06 | 7.29E-04 | 2.71 | 9950 | 132 | 668 | 24 |
| GO:1903901 | negative regulation of viral life cycle | 6.74E-06 | 7.71E-04 | 3.72 | 9950 | 60 | 668 | 15 |
| GO:0097529 | myeloid leukocyte migration | 6.99E-06 | 7.93E-04 | 3.38 | 9950 | 75 | 668 | 17 |
| GO:1901342 | regulation of vasculature development | 7.13E-06 | 7.96E-04 | 2.47 | 9950 | 169 | 668 | 28 |
| GO:0051480 | regulation of cytosolic calcium ion concentration | 7.37E-06 | 8.16E-04 | 2.62 | 9950 | 142 | 668 | 25 |

| | | | | | | | | |
|---|---|---|---|---|---|---|---|---|
| GO:0006816 | calcium ion transport | 7.56E-06 | 8.23E-04 | 2.82 | 9950 | 116 | 668 | 22 |
| GO:0048247 | lymphocyte chemotaxis | 7.53E-06 | 8.27E-04 | 5.32 | 9950 | 28 | 668 | 10 |
| GO:0007189 | adenylate cyclase-activating G protein-coupled receptor signaling pathway | 7.90E-06 | 8.34E-04 | 4.82 | 9950 | 34 | 668 | 11 |
| GO:0019932 | second-messenger-mediated signaling | 9.17E-06 | 9.53E-04 | 2.72 | 9950 | 126 | 668 | 23 |
| GO:1902105 | regulation of leukocyte differentiation | 1.02E-05 | 1.04E-03 | 2.34 | 9950 | 191 | 668 | 30 |
| GO:0019933 | cAMP-mediated signaling | 1.29E-05 | 1.29E-03 | 4.26 | 9950 | 42 | 668 | 12 |
| GO:0042113 | B cell activation | 1.35E-05 | 1.33E-03 | 2.89 | 9950 | 103 | 668 | 20 |
| GO:0006413 | translational initiation | 1.37E-05 | 1.34E-03 | 2.66 | 9950 | 129 | 668 | 23 |
| GO:0010469 | regulation of signaling receptor activity | 1.40E-05 | 1.36E-03 | 3.36 | 9950 | 71 | 668 | 16 |
| GO:0030217 | T cell differentiation | 1.46E-05 | 1.39E-03 | 3.08 | 9950 | 87 | 668 | 18 |
| GO:0051897 | positive regulation of protein kinase B signaling | 1.46E-05 | 1.40E-03 | 3.08 | 9950 | 87 | 668 | 18 |
| GO:0002690 | positive regulation of leukocyte chemotaxis | 1.70E-05 | 1.54E-03 | 3.66 | 9950 | 57 | 668 | 14 |
| GO:0019730 | antimicrobial humoral response | 1.69E-05 | 1.55E-03 | 4.16 | 9950 | 43 | 668 | 12 |
| GO:0002688 | regulation of leukocyte chemotaxis | 1.69E-05 | 1.55E-03 | 3.31 | 9950 | 72 | 668 | 16 |
| GO:0070374 | positive regulation of ERK1 and ERK2 cascade | 2.43E-05 | 2.18E-03 | 2.78 | 9950 | 107 | 668 | 20 |
| GO:0045765 | regulation of angiogenesis | 3.13E-05 | 2.67E-03 | 2.42 | 9950 | 154 | 668 | 25 |
| GO:0070838 | divalent metal ion transport | 3.24E-05 | 2.75E-03 | 2.47 | 9950 | 145 | 668 | 24 |
| GO:0042108 | positive regulation of cytokine biosynthetic process | 3.44E-05 | 2.91E-03 | 3.65 | 9950 | 53 | 668 | 13 |
| GO:1904018 | positive regulation of vasculature development | 3.52E-05 | 2.96E-03 | 2.8 | 9950 | 101 | 668 | 19 |
| GO:0072511 | divalent inorganic cation transport | 3.64E-05 | 3.03E-03 | 2.45 | 9950 | 146 | 668 | 24 |
| GO:0009164 | nucleoside catabolic process | 3.68E-05 | 3.05E-03 | 5.67 | 9950 | 21 | 668 | 8 |
| GO:0007187 | G protein-coupled receptor signaling pathway, coupled to cyclic nucleotide second messenger | 4.69E-05 | 3.79E-03 | 3.36 | 9950 | 62 | 668 | 14 |
| GO:0000956 | nuclear-transcribed mRNA catabolic process | 5.25E-05 | 4.20E-03 | 2.26 | 9950 | 178 | 668 | 27 |

| | | | | | | | | |
|---|---|---|---|---|---|---|---|---|
| GO:0045580 | regulation of T cell differentiation | 6.15E-05 | 4.80E-03 | 2.7 | 9950 | 105 | 668 | 19 |
| GO:2000106 | regulation of leukocyte apoptotic process | 6.81E-05 | 5.23E-03 | 3.26 | 9950 | 64 | 668 | 14 |
| GO:0000904 | cell morphogenesis involved in differentiation | 6.81E-05 | 5.26E-03 | 3.26 | 9950 | 64 | 668 | 14 |
| GO:0050870 | positive regulation of T cell activation | 7.88E-05 | 5.98E-03 | 2.34 | 9950 | 153 | 668 | 24 |
| GO:0043901 | negative regulation of multi-organism process | 7.97E-05 | 6.02E-03 | 2.5 | 9950 | 125 | 668 | 21 |
| GO:0042035 | regulation of cytokine biosynthetic process | 8.13E-05 | 6.11E-03 | 3.06 | 9950 | 73 | 668 | 15 |
| GO:0045766 | positive regulation of angiogenesis | 8.53E-05 | 6.37E-03 | 2.81 | 9950 | 90 | 668 | 17 |
| GO:0070663 | regulation of leukocyte proliferation | 8.76E-05 | 6.51E-03 | 2.32 | 9950 | 154 | 668 | 24 |
| GO:0007188 | adenylate cyclase-modulating G protein-coupled receptor signaling pathway | 9.45E-05 | 6.98E-03 | 3.34 | 9950 | 58 | 668 | 13 |
| GO:1903039 | positive regulation of leukocyte cell-cell adhesion | 1.00E-04 | 7.34E-03 | 2.26 | 9950 | 165 | 668 | 25 |
| GO:0050853 | B cell receptor signaling pathway | 1.04E-04 | 7.50E-03 | 4.47 | 9950 | 30 | 668 | 9 |
| GO:0019935 | cyclic-nucleotide-mediated signaling | 1.06E-04 | 7.59E-03 | 3.5 | 9950 | 51 | 668 | 12 |
| GO:0050663 | cytokine secretion | 1.15E-04 | 8.14E-03 | 4.03 | 9950 | 37 | 668 | 10 |
| GO:0022409 | positive regulation of cell-cell adhesion | 1.24E-04 | 8.63E-03 | 2.19 | 9950 | 177 | 668 | 26 |
| GO:0006412 | translation | 1.24E-04 | 8.68E-03 | 2.19 | 9950 | 177 | 668 | 26 |
| GO:0050679 | positive regulation of epithelial cell proliferation | 1.23E-04 | 8.69E-03 | 2.84 | 9950 | 84 | 668 | 16 |
| GO:0002443 | leukocyte mediated immunity | 1.37E-04 | 9.32E-03 | 3.23 | 9950 | 60 | 668 | 13 |
| GO:0002181 | cytoplasmic translation | 1.47E-04 | 9.83E-03 | 3.92 | 9950 | 38 | 668 | 10 |
| GO:0006402 | mRNA catabolic process | 1.78E-04 | 1.16E-02 | 2.11 | 9950 | 191 | 668 | 27 |
| GO:0051896 | regulation of protein kinase B signaling | 2.08E-04 | 1.35E-02 | 2.4 | 9950 | 124 | 668 | 20 |
| GO:0035456 | response to interferon-beta | 2.13E-04 | 1.38E-02 | 5.21 | 9950 | 20 | 668 | 7 |
| GO:0034341 | response to interferon-gamma | 2.39E-04 | 1.53E-02 | 2.79 | 9950 | 80 | 668 | 15 |
| GO:0046637 | regulation of alpha-beta T cell differentiation | 2.63E-04 | 1.62E-02 | 3.41 | 9950 | 48 | 668 | 11 |
| GO:0001764 | neuron migration | 2.63E-04 | 1.63E-02 | 3.41 | 9950 | 48 | 668 | 11 |

| | | | | | | | | |
|---|---|---|---|---|---|---|---|---|
| GO:0002449 | lymphocyte mediated immunity | 2.63E-04 | 1.64E-02 | 3.41 | 9950 | 48 | 668 | 11 |
| GO:2000257 | regulation of protein activation cascade | 2.80E-04 | 1.66E-02 | 4.41 | 9950 | 27 | 668 | 8 |
| GO:0000902 | cell morphogenesis | 2.81E-04 | 1.66E-02 | 2.48 | 9950 | 108 | 668 | 18 |
| GO:0030449 | regulation of complement activation | 2.80E-04 | 1.67E-02 | 4.41 | 9950 | 27 | 668 | 8 |
| GO:0034765 | regulation of ion transmembrane transport | 2.79E-04 | 1.68E-02 | 2.08 | 9950 | 186 | 668 | 26 |
| GO:0002920 | regulation of humoral immune response | 3.00E-04 | 1.77E-02 | 3.94 | 9950 | 34 | 668 | 9 |
| GO:0050670 | regulation of lymphocyte proliferation | 3.08E-04 | 1.80E-02 | 2.23 | 9950 | 147 | 668 | 22 |
| GO:0032944 | regulation of mononuclear cell proliferation | 3.40E-04 | 1.96E-02 | 2.21 | 9950 | 148 | 668 | 22 |
| GO:0050678 | regulation of epithelial cell proliferation | 3.43E-04 | 1.97E-02 | 2.17 | 9950 | 158 | 668 | 23 |
| GO:0032655 | regulation of interleukin-12 production | 3.58E-04 | 2.05E-02 | 3.55 | 9950 | 42 | 668 | 10 |
| GO:2000403 | positive regulation of lymphocyte migration | 3.69E-04 | 2.10E-02 | 4.26 | 9950 | 28 | 668 | 8 |
| GO:0070613 | regulation of protein processing | 3.72E-04 | 2.11E-02 | 2.93 | 9950 | 66 | 668 | 13 |
| GO:2000107 | negative regulation of leukocyte apoptotic process | 3.80E-04 | 2.14E-02 | 3.83 | 9950 | 35 | 668 | 9 |
| GO:0060337 | type I interferon signaling pathway | 3.85E-04 | 2.16E-02 | 3.28 | 9950 | 50 | 668 | 11 |
| GO:0043043 | peptide biosynthetic process | 3.92E-04 | 2.19E-02 | 2.04 | 9950 | 190 | 668 | 26 |
| GO:0050918 | positive chemotaxis | 4.17E-04 | 2.30E-02 | 4.74 | 9950 | 22 | 668 | 7 |
| GO:0007517 | muscle organ development | 4.39E-04 | 2.37E-02 | 3.46 | 9950 | 43 | 668 | 10 |
| GO:0032663 | regulation of interleukin-2 production | 4.39E-04 | 2.38E-02 | 3.46 | 9950 | 43 | 668 | 10 |
| GO:1903317 | regulation of protein maturation | 4.34E-04 | 2.38E-02 | 2.89 | 9950 | 67 | 668 | 13 |
| GO:0050671 | positive regulation of lymphocyte proliferation | 4.73E-04 | 2.50E-02 | 2.54 | 9950 | 94 | 668 | 16 |
| GO:1901658 | glycosyl compound catabolic process | 4.80E-04 | 2.52E-02 | 4.11 | 9950 | 29 | 668 | 8 |
| GO:0070372 | regulation of ERK1 and ERK2 cascade | 4.96E-04 | 2.55E-02 | 2.16 | 9950 | 152 | 668 | 22 |
| GO:0051607 | defense response to virus | 4.94E-04 | 2.57E-02 | 2.2 | 9950 | 142 | 668 | 21 |
| GO:0002793 | positive regulation of peptide secretion | 4.94E-04 | 2.57E-02 | 2.11 | 9950 | 162 | 668 | 23 |
| GO:0050731 | positive regulation of peptidyl-tyrosine phosphorylation | 5.35E-04 | 2.71E-02 | 2.51 | 9950 | 95 | 668 | 16 |

| | | | | | | | | |
|---|---|---|---|---|---|---|---|---|
| GO:0050715 | positive regulation of cytokine secretion | 5.40E-04 | 2.72E-02 | 2.6 | 9950 | 86 | 668 | 15 |
| GO:0032946 | positive regulation of mononuclear cell proliferation | 5.35E-04 | 2.72E-02 | 2.51 | 9950 | 95 | 668 | 16 |
| GO:0071347 | cellular response to interleukin-1 | 5.35E-04 | 2.73E-02 | 3.39 | 9950 | 44 | 668 | 10 |
| GO:0002460 | adaptive immune response based on somatic recombination of immune receptors built from immunoglobulin superfamily domains | 5.50E-04 | 2.76E-02 | 3.15 | 9950 | 52 | 668 | 11 |
| GO:0002548 | monocyte chemotaxis | 5.65E-04 | 2.81E-02 | 4.53 | 9950 | 23 | 668 | 7 |
| GO:0051282 | regulation of sequestering of calcium ion | 5.84E-04 | 2.89E-02 | 2.81 | 9950 | 69 | 668 | 13 |
| GO:0050714 | positive regulation of protein secretion | 5.96E-04 | 2.94E-02 | 2.13 | 9950 | 154 | 668 | 22 |
| GO:0046632 | alpha-beta T cell differentiation | 6.16E-04 | 3.00E-02 | 3.97 | 9950 | 30 | 668 | 8 |
| GO:0031341 | regulation of cell killing | 6.28E-04 | 3.05E-02 | 2.93 | 9950 | 61 | 668 | 12 |
| GO:0050851 | antigen receptor-mediated signaling pathway | 6.42E-04 | 3.11E-02 | 2.08 | 9950 | 165 | 668 | 23 |
| GO:2000401 | regulation of lymphocyte migration | 6.47E-04 | 3.12E-02 | 3.31 | 9950 | 45 | 668 | 10 |
| GO:0070665 | positive regulation of leukocyte proliferation | 6.78E-04 | 3.26E-02 | 2.46 | 9950 | 97 | 668 | 16 |
| GO:0045069 | regulation of viral genome replication | 6.95E-04 | 3.33E-02 | 2.64 | 9950 | 79 | 668 | 14 |
| GO:0001938 | positive regulation of endothelial cell proliferation | 7.78E-04 | 3.61E-02 | 3.24 | 9950 | 46 | 668 | 10 |
| GO:0050854 | regulation of antigen receptor-mediated signaling pathway | 7.71E-04 | 3.62E-02 | 3.03 | 9950 | 54 | 668 | 11 |
| GO:0001667 | ameboidal-type cell migration | 7.76E-04 | 3.62E-02 | 2.73 | 9950 | 71 | 668 | 13 |
| GO:0070588 | calcium ion transmembrane transport | 7.84E-04 | 3.63E-02 | 2.51 | 9950 | 89 | 668 | 15 |
| GO:0010951 | negative regulation of endopeptidase activity | 8.42E-04 | 3.86E-02 | 2.27 | 9950 | 118 | 668 | 18 |
| GO:0030183 | B cell differentiation | 8.50E-04 | 3.87E-02 | 2.84 | 9950 | 63 | 668 | 12 |
| GO:0042476 | odontogenesis | 8.97E-04 | 4.05E-02 | 3.44 | 9950 | 39 | 668 | 9 |
| GO:0070848 | response to growth factor | 8.99E-04 | 4.05E-02 | 2.03 | 9950 | 169 | 668 | 23 |
| GO:0010466 | negative regulation of peptidase activity | 9.32E-04 | 4.15E-02 | 2.25 | 9950 | 119 | 668 | 18 |
| GO:0061061 | muscle structure development | 9.30E-04 | 4.16E-02 | 3.17 | 9950 | 47 | 668 | 10 |

| | | | | | | | | |
|---|---|---|---|---|---|---|---|---|
| GO:0001936 | regulation of endothelial cell proliferation | 9.84E-04 | 4.30E-02 | 2.79 | 9950 | 64 | 668 | 12 |
| GO:0034656 | nucleobase-containing small molecule catabolic process | 9.82E-04 | 4.32E-02 | 3.72 | 9950 | 32 | 668 | 8 |
| GO:0032649 | regulation of interferon-gamma production | 9.84E-04 | 4.32E-02 | 2.79 | 9950 | 64 | 668 | 12 |

### Supplementary Table 4: Hospital cohort clinical indications information

| Patient ID | HIV status | Prior TB | Clinical indication | Final Diagnosis |
|---|---|---|---|---|
| 012-09-0106 | positive | No | Exclude infection | No active infectious or inflammatory disease |
| 012-09-1009 | positive | No | Exclude infection | No active infectious or inflammatory disease |
| 012-09-1040 | positive | Yes | Exclude infection | No active infectious or inflammatory disease |
| 012-09-1043 | negative | No | Exclude infection/Malignancy | No active infectious or inflammatory disease |
| 012-09-1049 | negative | No | Exclude TB | No active infectious or inflammatory disease |
| 012-09-1058 | negative | No | Exclude infection | No active infectious or inflammatory disease |
| 012-09-1027 | negative | No | Exclude infection | No active infectious or inflammatory disease |
